# Pharmacological Antagonism of Ccr2+ Cell Recruitment to Facilitate Regenerative Tendon Healing

**DOI:** 10.1101/2024.07.15.603448

**Authors:** Gilbert Smolyak, Andrew Rodenhouse, Anne E.C. Nichols, Constantinos Ketonis, Alayna E. Loiselle

## Abstract

Successful tendon healing requires sufficient deposition and remodeling of new extracellular matrix at the site of injury, with this process mediating in part through fibroblast activation via communication with macrophages. Moreover, resolution of healing requires clearance or reversion of activated cells, with chronic interactions with persistent macrophages impairing resolution and facilitating the conversion the conversion to fibrotic healing. As such, modulation of the macrophage environment represents an important translational target to improve the tendon healing process. Circulating monocytes are recruited to sites of tissue injury, including the tendon, via upregulation of cytokines including Ccl2, which facilitates recruitment of Ccr2+ macrophages to the healing tendon. Our prior work has demonstrated that Ccr2-/-can modulate fibroblast activation and myofibroblast differentiation. However, this approach lacked temporal control and resulted in healing impairments. Thus, in the current study we have leveraged a Ccr2 antagonist to blunt macrophage recruitment to the healing tendon in a time-dependent manner. We first tested the effects of Ccr2 antagonism during the acute inflammatory phase and found that this had no effect on the healing process. In contrast, Ccr2 antagonism during the late inflammatory/ early proliferative period resulted in significant improvements in mechanical properties of the healing tendon. Collectively, these data demonstrate the temporally distinct impacts of modulating Ccr2+ cell recruitment and Ccr2 antagonism during tendon healing and highlight the translational potential of transient Ccr2 antagonism to improve the tendon healing process.

## INTRODUCTION

Tendons allow the mechanical transmission of force from muscles to bones facilitating movement across the entire skeleton. Injuries to tendons are very common, comprising about half of all musculoskeletal injuries every year.^1^ Surgical intervention is typically required to repair acute, traumatic tendon transections. However, there are several complications associated with these procedures including: joint stiffness, re-rupture of the repaired tendon, infection, and most commonly peritendinous adhesion formation.^2, 3^ Adhesions restrict range of motion by adhering tendon to surrounding structures due to an exuberant extracellular matrix (ECM) response to injury.^4^

While the precise drivers of the fibrotic response to tendon injury are incompletely defined, immune cells, particularly macrophages are key regulators of both physiologic and fibrotic healing.^5^ During the acute inflammatory phase of healing, circulating monocytes/macrophages are recruited to the healing tendon via chemokine signals, including CCL2 (*Mcp*-1, monocyte chemoattractant protein-1),^6, 7^ and subsequently aid in propagating the inflammatory response which facilitates recruitment of other cell populations and increases vascularity^8^. Following resolution of acute inflammation, some macrophages remain in the healing tendon and may contribute to fibroblast activation, as observed in other tissues^9^, which can facilitate extracellular matrix (ECM) production and remodeling.^9–11^ While production of a provisional matrix is a key step in healing, chronic macrophage-fibroblast interactions can facilitate a switch from physiologic to fibrotic wound healing^12–14^, with matrix production outpacing remodeling.^11^ Thus, macrophages are an attractive candidate for therapeutic modulation as a means to improve the tendon healing process. We have previously used a Ccr2^-/-^ model, which blunts recruitment of circulating monocytes/macrophages to the site of injury to blunt macrophage recruitment to the healing tendon, however this approach does not allow for any temporal control of modulating macrophage recruitment, and Ccr2^-/-^ resulted in healing impairments ^15^. In contrast, mild attenuation of macrophage presence has shown some beneficial effects on the healing process.^16–18^ Collectively, these data suggest that a more translationally-tenable therapeutic approach to enhancing healing via modulation of the macrophage environment is likely to involve blunting rather than complete attenuation of macrophage recruitment and temporal control of macrophage dynamics. To that end, we leveraged a small molecule antagonist of Ccr2, which blunts circulating monocyte/macrophage recruitment to sites of injury or inflammation,^19^ and hypothesized that reducing macrophage recruitment to the healing tendon following the acute inflammatory period would enhance the tendon healing process.

## METHODS

### Animal Ethics and Husbandry

This study was carried out in strict accordance with the recommendations in the Guide for the Care and Use of Laboratory Animals of the National Institutes of Health. All animal procedures were approved by the University Committee on Animal Research (UCAR) at the University of Rochester (UCAR Number: 2014-004E). Mice were housed with up to five animals per cage with *ab libitum* access to water and standard chow, on a 12:12 light: dark cycle. Animals were randomly assigned to experimental and control groups, and all animals that were enrolled in the study completed the study and were sacrificed at the planned timepoint. No adverse events occurred.

### Tendon Repair Surgery

Mice underwent complete transection and repair of the flexor digitorum longus (FDL) tendon in the hind paw as previously described.^20–25^ Briefly, female C57Bl/6J (#664, Jackson Laboratories) mice aged 10-12 weeks were sedated with Ketamine/Xylazine and received pre-operative analgesic via extended-release buprenorphine. Following preparation of the surgical site, an incision was made on medial aspect of the right lower limb and the myotendinous junction was identified and transected. The skin was closed using 5-0 nylon suture. Another incision was made on the plantar aspect of the right foot and the distal portion of the FDL tendon was identified, fully transected, and subsequently repaired with an 8-0 nylon suture using a modified Kessler repair technique. Skin was closed with a 5-0 nylon suture.

### Ccr2 Antagonist Treatment

Mice were treated with 5mg/kg of the Ccr2 antagonist, RS102895,^19^ every 8hrs during the treatment windows outlined in Figures 1A, 2A, and 4A. Vehicle control mice were treated with a 60:40 mix of DMSO:saline. Schematics were created using BioRender.com.

### Histology and Immunofluorescence

Tissues (n=4-5 per group per treatment regimen) were harvested on post-operative day 14, fixed in 10% neutral buffer formalin for 72h at room temperature, decalcified in Webb-Jee 14% EDTA solution for 14d, processed, and embedded in paraffin. Three-micron sections were cut through the transverse plane of the hind paw. Slides were stained with either Alcian Blue Hematoxylin/ Orange G, Masson’s Trichrome, or used for immunostaining. For immunofluorescent staining, tissue sections were probed with either an anti-Ccr2 primary (1:100, NOVUS SN707) and FITC-conjugated Donkey-anti-rabbit secondary antibody (1:200, Jackson ImmunoResearch, #711-096-152, RRID: AB_2340597), or an anti-αSMA-Cy3 antibody (1:250, Sigma-Aldrich, #C6198). Nuclei were counterstained with NucBlue Live Cell Stain (Invitrogen, #R37605). Slides were imaged with a VS120 Virtual Slide Microscope. Immunostaining was quantified in a semi-automated manner using Visiopharm (Version 2022.01.3.12053) as we have previously described.^15^ The percent positive area of αSMA was calculated within a consistent region of interest based on an input threshold, while the percent of Ccr2+ cells was calculated as a fraction of total cells (number of nuclei) within the region of interest.

### Range of motion assessment and biomechanical testing

After harvest on post-operative day 14, hind paws were prepared for range of motion (ROM) analyses as previously described.^26, 27^ Briefly, the proximal FDL tendon was freed from surrounding connective tissue, and secured between two pieces of tape with cyanoacrylate. The tibias of these samples were then fastened with in clip such that the plantar aspect of the foot was pointed towards the celling and weights of increasing weights (1, 3, 5, 8, 12, and 19g) were incrementally applied to the tendon, and the resulting flexion angle at the MTP joint was recorded for each weight using a digital camera and then measured using ImageJ (version 1.54f). Gliding resistance was calculated using the change in flexion angles over the range of applied loads as previously described^27^ with a higher gliding resistance indicated an impaired ability of the tendon to move freely, and as is associated with peritendinous adhesion formation.

Samples were then prepared for mechanical analysis by removing the tibia at the ankle and freeing the tendon from the tarsal tunnel. The samples were fastened in custom grips mounted on an Instron ELECTROPULS® E10000 Linear-Torsion all-electric dynamic test instrument, with one grip securing the distal aspect of the foot and the other securing the proximal tendon. A linear axial tensile force was then applied at a rate of 30mm/min until tendon failure. Maximum load at failure, and stiffness (slope of the linear region of the force-displacement curve) were then calculated. Twelve animals per group per treatment regimen were used. Samples that were damaged during harvest or testing prep were excluded.

### Statistical Analysis

Quantitative data were analyzed using GraphPad Prism (version 10.2.2 (397)). Outliers were identified using the ROUT method (Q value = 1%). A student’s t-test was used to identify statistically significant differences between control and experimental groups. Sample sizes were determined from post-hoc power calculations of our prior studies. For all experiments, n = 1 represents one mouse. For all statistical analyses p < 0.05 was considered to be statistically significant. Significance is reported in all figures with the following conventions: * = p<0.05, ** = p<0.01, *** = p<0.001, **** = p<0.0001.

## RESULTS

### RS102895 blunts recruitment of Ccr2+ cells to the healing tendon

To confirm that the Ccr2 antagonist, RS102895 was sufficient to blunt Ccr2+ cell recruitment during tendon healing, mice (n=3 per group) were treated from day 0-3 and tendons were harvested at day 5 (Figure 1A). A substantial decline in the proportion of Ccr2+ cells was observed in antagonist treated mice (RS102985: 42.42 ± 11.85%; Vehicle: 60.99 ± 7.144%), although this decline was not significantly different (p = 0.0808), likely due to the small sample size in this pilot experiment (Figure 1B and 1C). Based on these promising results, we tested the functional and mechanical implications of RS102895 treatment during two different treatment windows.

**Figure 1.**
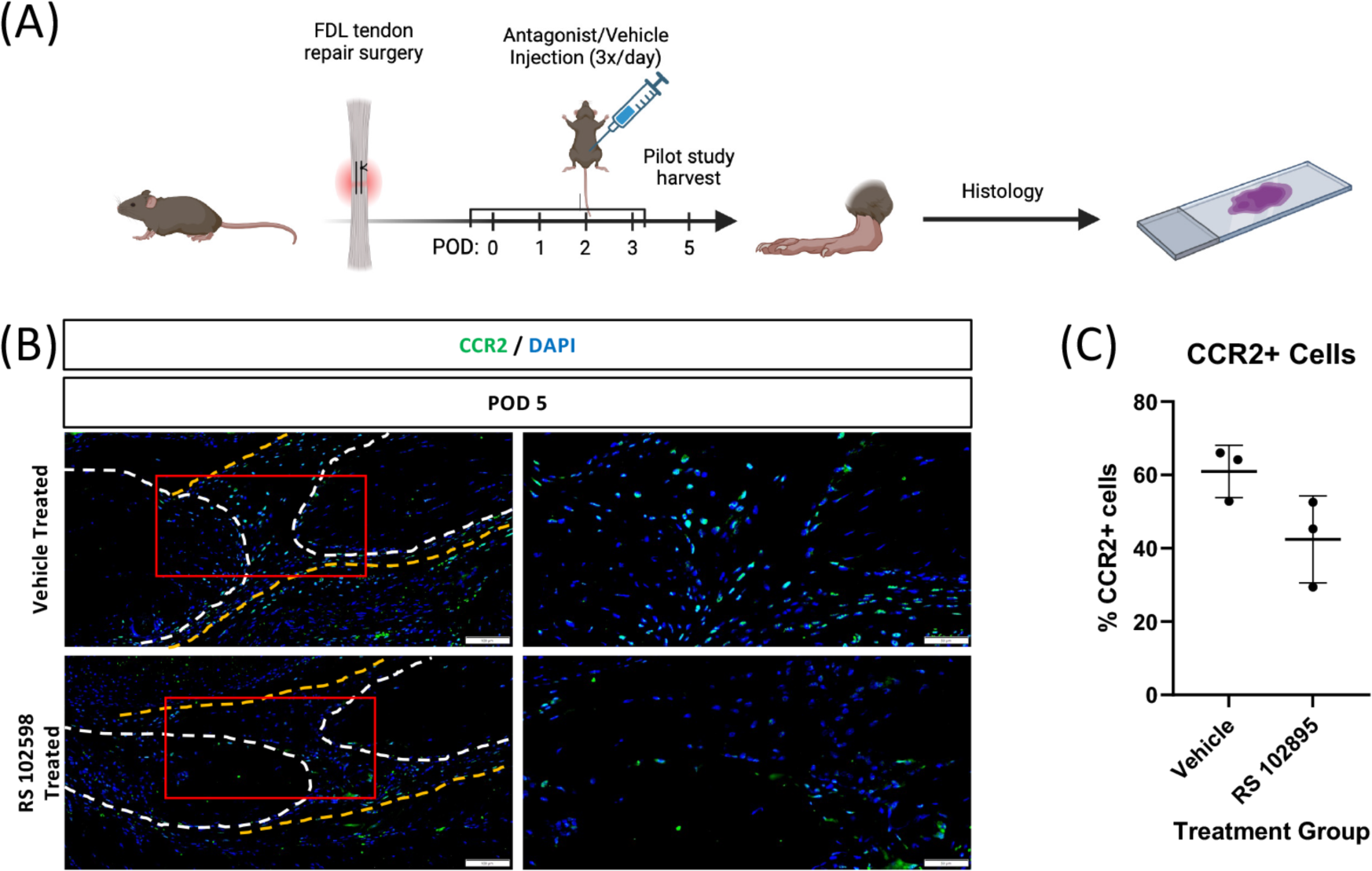
(A) Schematic of pilot experiment: Mice were injected with a CCR2 antagonist (RS102895) every 8hrs beginning just prior to surgery, with treatment through day 3, and healing tendons harvested at post-operative day (POD) 5 to assess for efficacy of Ccr2 antagonism via quantification of Ccr2+ cells. (B) Immunofluorescence results from the pilot study with stubs of the native tendon outlined in white and bridging tissue outlined in orange. Red box indicates area of higher magnification image. (C) Quantification of the percent of Ccr2+ cells in the healing tendon.

### Ccr2 inhibition during acute inflammation (day 0-4) does not alter healing

To determine the effects of Ccr2 inhibition in the early inflammatory phase on functional and mechanical recovery, mice were treated with a Ccr2 antagonist, or vehicle control, from day 0-4 to target the initial influx of circulating monocytes/macrophages, and tissues were harvested at post-operative day (POD) 14 (Figure 2A). No significant differences were found between vehicle and RS 102895 treated mice in terms of gliding resistance (Vehicle: 32.43 ± 13.20; RS102895: 27.70 ± 6.84, p = 0.29), MTP flexion angle (Vehicle: 29.37° ± 13.26; RS102895: 32.53° ± 5.24, p = 0.45), tendon stiffness (Vehicle: 2.49N/mm ± 1.35; RS102895: 2.55N/mm ± 1.67, p = 0.92), and maximum load at failure (Vehicle: 0.98N ± 0.55; RS102895: 0.88N ± 0.19, p=0.59) (Figure 2B-D), suggesting that pharmacological antagonism of Ccr2 in the early inflammatory phase does not substantially alter functional and mechanical restoration during tendon healing.

We then further interrogated whether there were changes in tissue morphology and composition of the cell environment. No significant difference in the percent of Ccr2+ cells was observed between vehicle and antagonist treated repairs (Vehicle: 40.93 ± 26.58; RS102985: 26.85 ± 16.49, p = 0.3882) (Figure 3A & B), likely due to a subsequent influx of Ccr2+ after the cessation of antagonist treatment. Further, there was no significant difference in the area of αSMA staining between the vehicle and antagonist treated mice (16.10 ± 11.25% vs 21.33 ± 7.659% respectively, p = 0.4149) (Figure 3C & D). In addition, there were no observable morphological differences between vehicle and antagonist treated tendons using both ABHOG (Figure 3E) and Masson’s Trichrome staining (Figure 3F). Collectively, these data suggest that Ccr2 antagonism to temper the recruitment of macrophages during acute inflammation does not have lasting effects on the healing environment at 14 days after injury.

**Figure 2.**
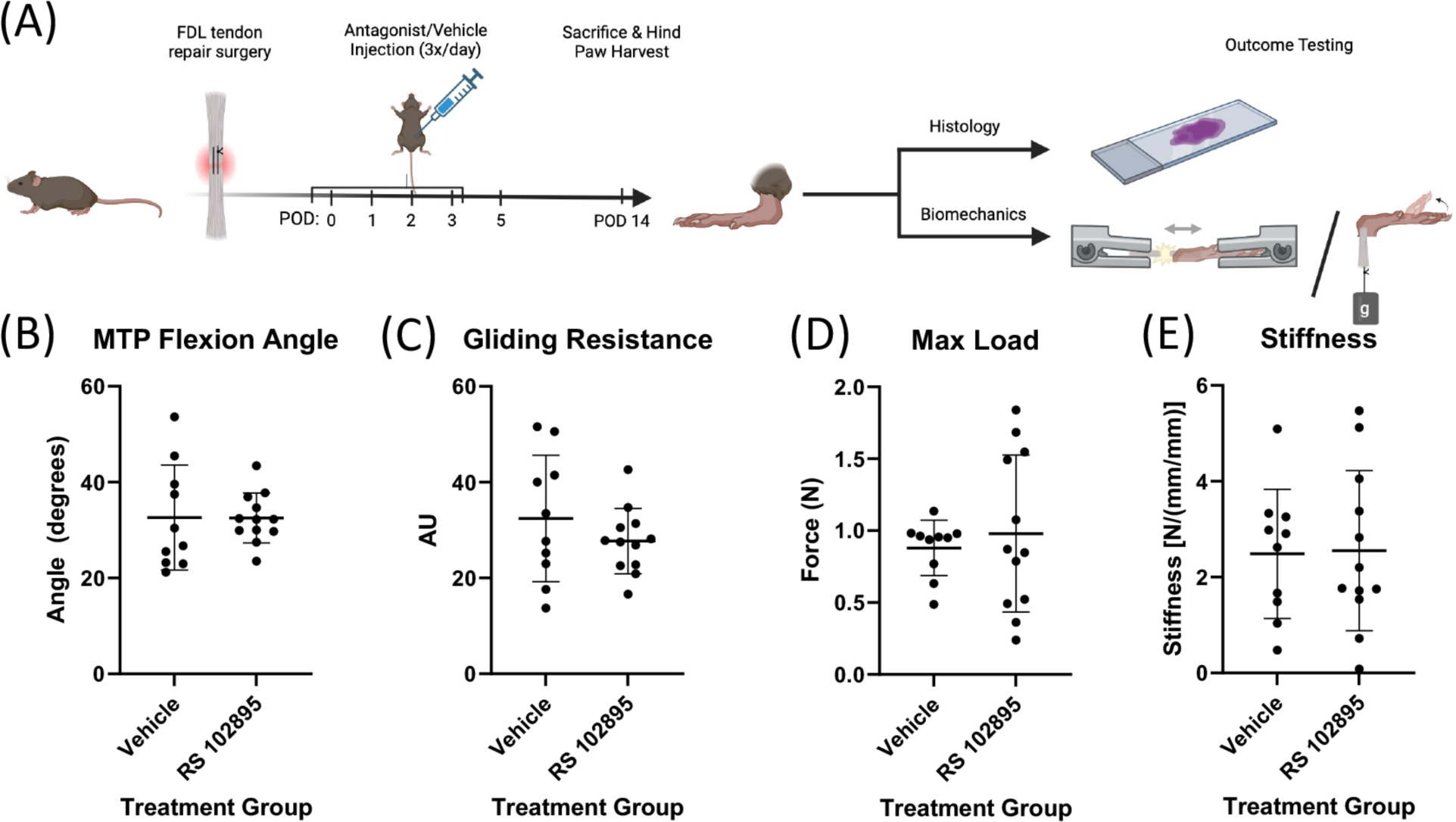
(A) Schematic of Ccr2 antagonism during the acute inflammatory phase (treatment from post-operative day (POD) 0-3 with tissue harvests at POD 14 for histological, biomechanical, and functional outcomes. (B) MTP Flexion Angle, (C) Gliding Resistance, (D) Max Load at Failure, and (E) Stiffness of healing tendons from vehicle and Ccr2 antagonist (RS102895) treated mice at POD 14.

**Figure 3.**
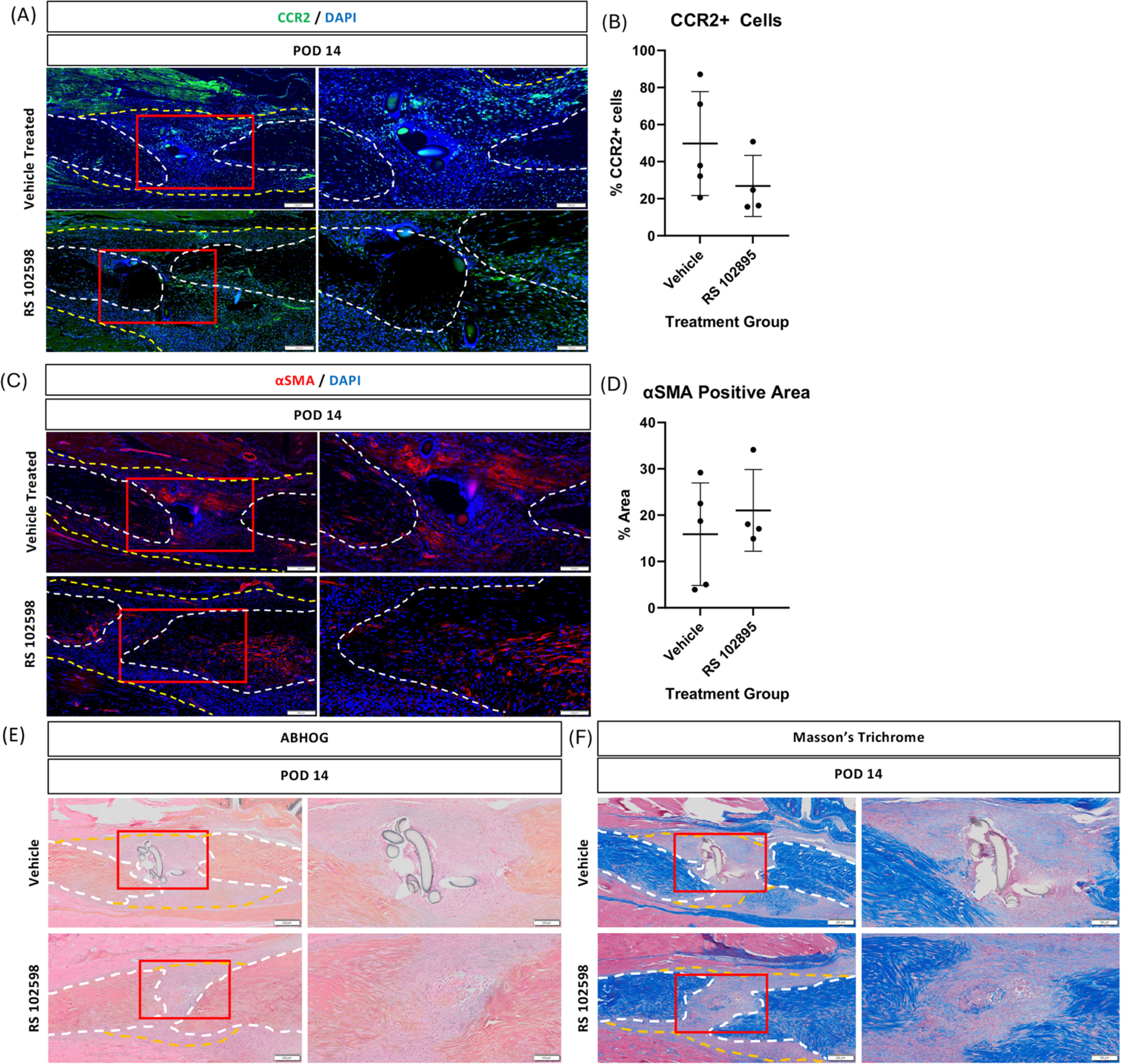
(A) Ccr2 immunostaining, (B) quantification of Ccr2+ cells, (C) αSMA immunostaining, (D) quantification of the αSMA+ area, (E) Alcian Blue/ Hematoxylin/ Orange G (ABHOG) staining, and (F) Masson’s Trichrome staining of healing tendons from mice treated with vehicle and Ccr2 antagonist (RS102985) from post-operative day (POD) 0-3, with tendons harvested at POD 14. Red box indicates area of higher magnification image.

### Ccr2 inhibition during the early proliferative phase (day 5-8) enhances healing

To determine the effect of Ccr2 antagonism in the early proliferative phase (day 5-8) on the healing process, healing tendons were harvested at POD 14 (Figure 4A). No significant differences in MTP flexion angle (p=0.78) and gliding resistance (p=0.61) were observed between vehicle control and antagonist treated repairs (Figure 4B and 4C). However, Ccr2 antagonist treated tendons demonstrated a significant increase in max load to failure (vehicle: 0.58 ± 0.16 N; RS102895: 0.82 ± 0.15 N, p = 0.0038) (Figure 4D), and stiffness (Vehicle: 1.85 ± 0.88 N/mm; RS102895: 2.76 ± 0.54 N/mm, p = 0.02) (Figure 4E), relative to vehicle treated repairs. Collectively, these data suggest that Ccr2 antagonism in the late inflammatory/early proliferative phase enhances the healing process at 14 days after injury.

We then began to define morphological and changes in the composition of the cell environment that are associated with enhanced healing. Consistent with the early RS102895 treatment regimen, there was no difference between the percent Ccr2+ cells in vehicle vs. antagonist treated repairs (Vehicle: 18.23 ± 19.37; RS102895: 25.59 ± 16.54 respectively, p = 0.54) (Figure 5A & B). Interestingly, while there was a qualitative increase in αSMA staining in vehicle treated mice, relative to antagonist treated, these differences were not significant (Vehicle: 14.25 ± 8.51%; RS102895: 7.13 ± 4.06%, p = 0.14, Figure 5C &D). ABHOG and Masson’s trichrome staining identified a consistent qualitative increase in the extent of degradation/remodeling of the native tendon stubs in RS102895 treated tendons (blue arrows, Figure 5E & F). Collectively, these results suggest that Ccr2 antagonism treatment at the beginning of the proliferative phase may improve healing via altered collagen deposition and remodeling.

**Figure 4.**
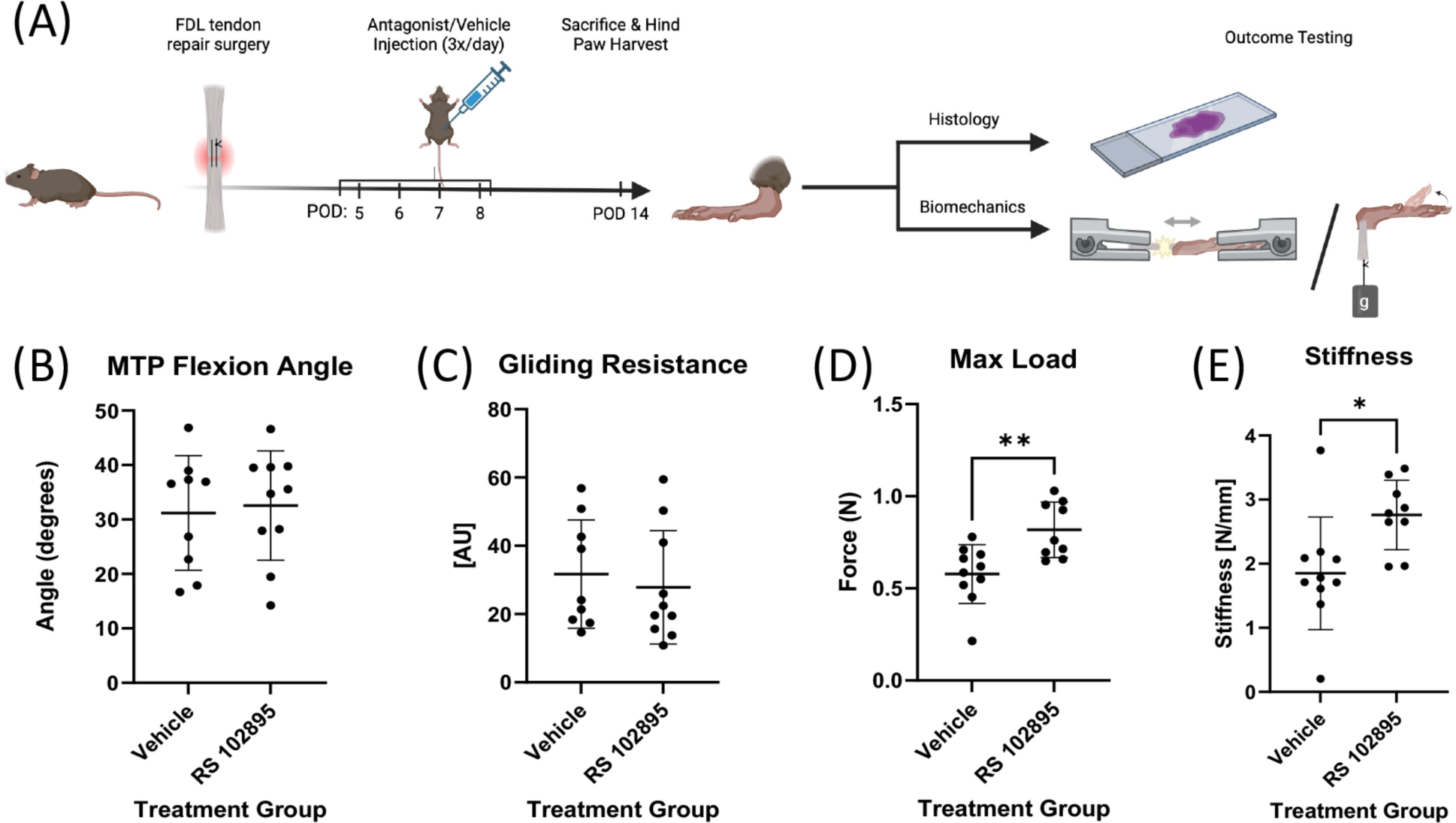
(A) Schematic of Ccr2 antagonism during the early proliferative phase (treatment from post-operative day (POD) 5-8 with tissue harvests at POD 14 for histological, biomechanical, and functional outcomes. (B) MTP Flexion Angle, (C) Gliding Resistance, (D) Max Load at Failure, and (E) Stiffness of healing tendons from vehicle and Ccr2 antagonist (RS102895) treated mice at POD 14.

**Figure 5.**
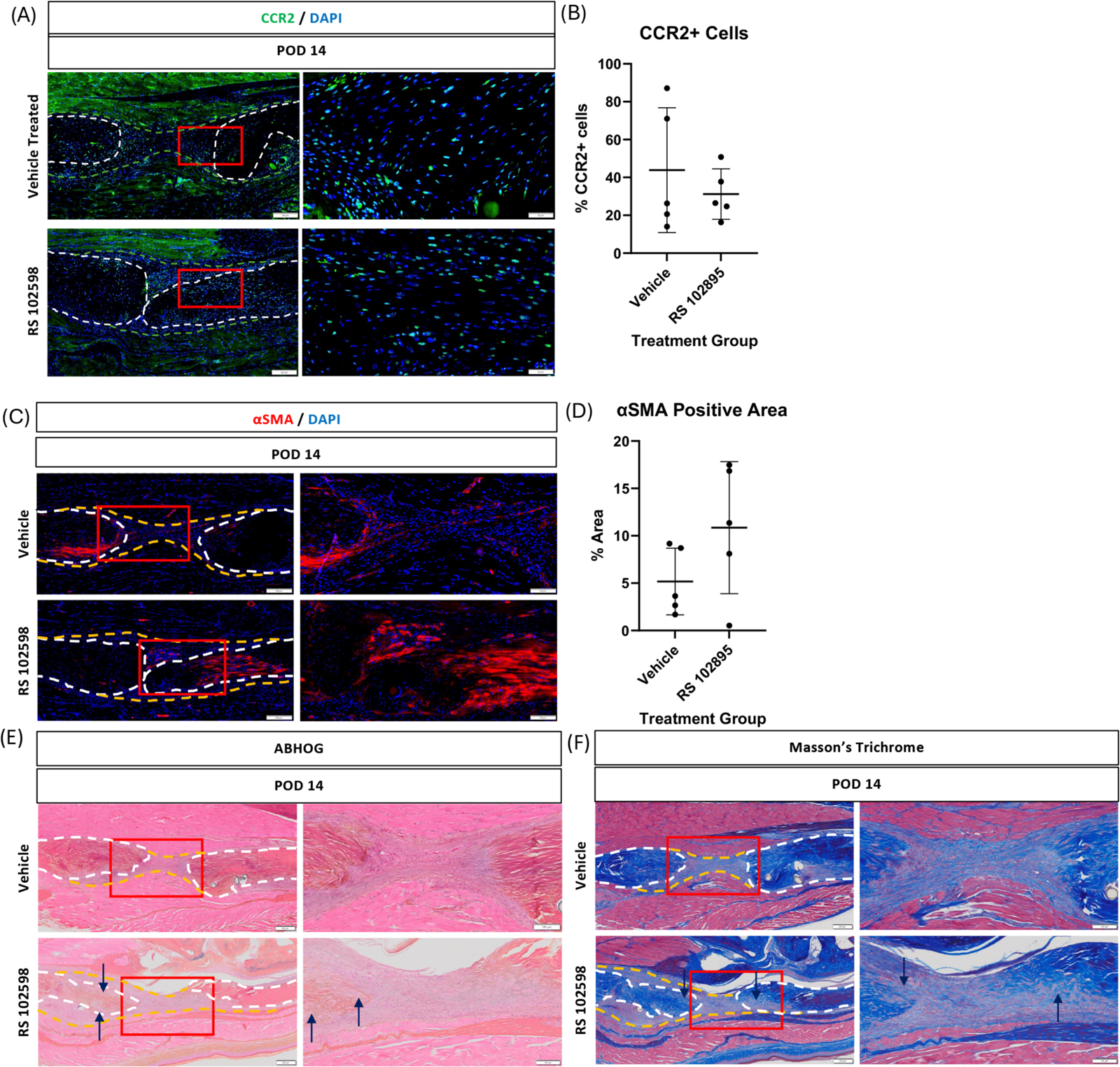
A) Ccr2 immunostaining, (B) quantification of Ccr2+ cells, (C) αSMA immunostaining, (D) quantification of the αSMA+ area, (E) Alcian Blue/ Hematoxylin/ Orange G (ABHOG) staining, and (F) Masson’s Trichrome staining of healing tendons from mice treated with vehicle and Ccr2 antagonist (RS102985) from post-operative day (POD) 5-8, with tendons harvested at POD 14. Red box indicates area of higher magnification image. Blue arrows indicate remodeling of the native tendon stubs.

## DISCUSSION

In the present study, we examined the effects of temporal antagonism of the Ccr2 receptor, with the goal of blunting macrophage infiltration into the healing tendon during distinct phases of healing (acute inflammatory vs. early proliferative phase) to establish the potentially divergent impacts on the healing process. While we have previously demonstrated that Ccr2^-/-^ impairs the healing process,^15^ the current study focuses on a more translational pharmacologic approach that allows temporal control of Ccr2-mediated macrophage recruitment. Given that macrophages can play both physiological (e.g., clearing debris from the wound area and activating surrounding cells) and pathological roles during healing (e.g., facilitating tissue fibrosis via chronic interactions with fibroblasts ^29^), we tested the hypothesis that Ccr2 antagonism in the peri-inflammatory period would reduce macrophage presence at the injury site resulting in improved functional outcomes. Indeed, Ccr2 antagonism from day 5-8 post-surgery significantly improved mechanical properties of healing tendons, relative to vehicle. In contrast, Ccr2 antagonism during the acute inflammatory phase did not markedly alter the healing process. Collectively, these data highlight the time-dependent functions of Ccr2 in the tendon healing process and identify an important therapeutic window for Ccr2 modulation to enhance healing.

While prior work has demonstrated that inhibition of the early inflammatory phase blunts initiation of the physiological healing cascade and impairs healing,^7, 15, 30, 31^ the effects of modulating the macrophage environment have demonstrated conflicting effects. For example, clodronate depletion of macrophages reduced extracellular matrix production while also enhancing reacquisition of mechanical properties.^17^ In contrast, Howell *et al*., demonstrated that macrophage depletion impairs healing of neonatal tendons.^14^ Moreover, while we have demonstrated that Ccr2^-/-^ impairs tendon healing,^15^ recent work has demonstrated that Ccr2^-/-^ enhances healing in a model of delayed repair of rotator cuff injuries,^32^ highlighting the context-dependent potential of modulating the macrophage environment. Moreover, the differences observed between Ccr2 antagonism from day 0-3 and day 5-8 highlight the temporally dependent nature of macrophage functions, particularly those mediated via Ccr2.

The efficacy of Ccr2 antagonism to blunt monocyte recruitment is well established including as a means to improve responsiveness to vaccine,^19^ decrease the burden of pancreatic cancer,^33^ improve outcomes after myocardial infarction,^34^ decrease renal fibrosis,^35^ and slow the progression of osteoarthritis.^36^ However, Ccr2 antagonism may modulate cell functions in addition to, or independent from, the effects on Ccr2 cell recruitment. For example, prior work suggests that RS102895 can promote pro-inflammatory macrophage polarization *in vitro*.^37^ However, this effect has not been observed in vivo, and the relative impacts of blunting macrophage recruitment, while promoting inflammatory functions of those macrophages that are able to respond to injury would be an important area for future investigation. Given that our focus was on disrupting macrophage recruitment in a time-dependent manner, we did not assess for changes in macrophage polarization, but rather examined how reduced Ccr2+ cell presence altered other aspects of the cell environment, with a particular focus on myofibroblasts. Interestingly, while our decision to treat with RS102895 during the late inflammatory/early proliferative phase was premised on preventing the chronic macrophage-myofibroblast interactions that are associated with a conversion to tissue fibrosis,^38^ we saw a qualitative, but non-significant increase in αSMA+ myofibroblasts, suggesting that the improvements in healing observed with Ccr2 antagonism are likely not directly the result of decreased myofibroblast presence. Future work to define broad changes in the cell environment with Ccr2 antagonism and blunted macrophage recruitment are needed to provide a clearer picture of the mechanisms through which Ccr2 antagonism from day 5-8 enhances healing, and to what extent macrophage-fibroblast/myofibroblast communication patterns are altered.

Despite the beneficial effects of delayed Ccr2 antagonism, there are limitations to our study that must be considered. First, only female mice were used in this experiment. While we have not previously observed substantial sexual dimorphism during healing in this murine model, it is possible that differences in antagonist metabolism may alter the treatment effect between males and females. In addition, we only examined the impact of antagonist treatment on the healing process with outcome measures at day 14 after surgery. Given that we have previously demonstrated that Ccr2^-/-^ impairs healing at 28 days, understanding whether the improvements in healing with Ccr2 antagonist treatment are maintained long-term is an important area for future studies. Finally, it is likely that we did not achieve consistent blunting of Ccr2 during the entirety of the treatment windows as RS102895 has a relatively short half-life, with treatment every 6hrs previously reported to be required to achieve consistent therapeutic dosing in vivo.^19^ However, we treated every 8hrs in an attempt to balance consistent Ccr2 antagonism with reduced post-operative animal handling which we have anecdotally observed can contribute to an increase in repair ruptures. As such, the impact of this treatment regimen may underestimate the beneficial effects of consistent Ccr2 antagonism.

Overall, this study established the differential effects of Ccr2 pharmacological antagonism during the acute inflammatory vs. early proliferative phase of healing and provides insight into the temporally dependent functions of Ccr2+ macrophages. Moreover, the improvements in restoration of tendon biomechanical properties with Ccr2 antagonism from day 5-8 after injury highlights the therapeutic potential of modulating Ccr2 and future studies can build on these exciting data to both define the mechanisms of enhanced healing and further refine the treatment window (e.g., dose, timing, duration) to establish a translationally feasible means of improving clinical tendon injuries.

